# Thalamic Network Controllability Predicts Cognitive Impairment in Multiple Sclerosis

**DOI:** 10.1101/2023.11.29.568586

**Authors:** Yuping Yang, Anna Woollams, Ilona Lipp, Zhizheng Zhuo, Marta Czime Litwińczuk, Valentina Tomassini, Yaou Liu, Nelson J Trujillo-Barreto, Nils Muhlert

## Abstract

Recent research suggests that individuals with multiple sclerosis (MS) and cognitive impairment exhibit more effortful and less efficient transitions in brain network activity. Previous studies further highlight the increased vulnerability of specific regions, particularly the thalamus, to disease-related damage. This study investigates whether MS affects the controllability of specific brain regions in driving network activity transitions across the brain and examines the relationship between these changes and cognitive impairment in patients. Resting-state functional MRI and neuropsychological data were collected from 102 MS and 27 healthy controls. Functional network controllability analysis was performed to quantify how specific regions influence transitions between brain activity patterns or states. Disease alterations in controllability were assessed in the main dataset and then replicated in an independent dataset of 95 MS and 45 healthy controls. Controllability metrics were then used to distinguish MS from healthy controls and predict cognitive status. MS-specific controllability changes were observed in the subcortical network, particularly the thalamus, which were further confirmed in the replication dataset. Cognitively impaired patients showed significantly greater difficulty in the thalamus steering brain transitions towards difficult-to-reach states, which are typically associated with high-energy-cost cognitive functions. Thalamic network controllability proved more effective than thalamic volume in distinguishing MS from healthy controls (AUC = 88.3%), and in predicting cognitive status in MS (AUC = 80.7%). This study builds on previous research highlighting early thalamic damage in MS, aiming to demonstrate how this damage disrupts activity transitions across the cerebrum and may predict cognitive deficits. Our findings suggest that the thalamus in MS becomes less capable of facilitating broader brain activity transitions essential for high-energy-cost cognitive functions, implying a potential pathological mechanism that links thalamic functional changes to cognitive impairment in MS.

## Introduction

Cognitive impairment is common in multiple sclerosis (MS), with estimated prevalence ranging from 43% to 70%.^1^ Previous studies suggest that cognitive impairments in MS arise due to brain changes in specific neural networks.^2–10^ A useful recent model indicates that MS patients with cognitive impairment (CIMS), in comparison to cognitively preserved patients (CPMS), exhibit less frequent transitions between brain states (regional activity patterns) and show subcortical-related functional connectivity changes.^5^ Subcortical network has gained wide attention in MS, with many studies reporting associations between cognitive impairment and network changes in subcortical regions, such as the thalamus.^6–8,10^ Central to these findings is the idea that subcortical regions, particularly the thalamus, are affected early in the course of MS;^11–15^ as the disease progresses, neurodegeneration and functional network changes seem to spread from the thalamus towards the broader brain.^16^ This early involvement of the thalamus may drive the subsequent changes in the other parts of the brain, leading to a widespread disruption of brain activity and causing reduced efficiency in brain function.

An emerging approach that allows characterizing how specific brain regions influence network activity in the rest of the brain is ‘network controllability’ analysis.^17–19^ This approach allows moving beyond assessing changes in regional activity, towards identifying the brain regions driving those changes and how they influence the transitions between brain states (regional activity patterns of the brain). Network controllability has been validated as a powerful tool in exploring clinical biomarkers in neurological and neuropsychological diseases.^20–24^ In MS, a recent study showed that it became more effortful for cognitively impaired patients to transition between brain states.^25^ However, current knowledge has not yet addressed whether and how MS alters the controllability of specific brain regions in facilitating transitions across the brain, and how these alterations predict cognitive impairment in patients.

To fill this gap, this study applied widely used controllability measures to two independent MS datasets to examine how different regions impact brain state transitions. These measures were further evaluated as predictors of cognitive impairment in MS and for their potential to support clinical diagnosis. We hypothesize that: (i) MS patients exhibit regions less effective in facilitating brain state transitions, as reflected in disease-related controllability changes; (ii) these changes are concentrated in those brain regions associated with cognitive dysfunction in MS, such as the thalamus; (iii) are more pronounced in cognitively impaired patients; and (iv) can help predict cognitive impairment and aid clinical diagnosis.

## Materials and Methods

### Participants

#### Main dataset

The main dataset involved 102 patients with relapsing-remitting MS recruited from the Helen Durham Centre for Neuroinflammation at the University Hospital of Wales and 27 healthy controls (HC) recruited from the local community. All participants were aged between 18 and 60 years, right-handed, and devoid of contraindications for MR scanning. Additional eligibility criteria were required for the patients including the absence of comorbid neurological or psychiatric disease, no modifications to their treatments within three months prior to the MRI scanning, and being in a relapse-free phase. All participants underwent demographic information collection, clinical and psychological assessments, and MRI scanning in one study session. This study was approved by the NHS South-West Ethics and the Cardiff and Vale University Health Board R&D committees. Written informed consent was obtained from each participant.

#### Replication dataset

The replication dataset included 95 patients with relapsing-remitting MS recruited from the Beijing Tiantan Hospital and 45 HC recruited from the local community. The participants were aged between 17 and 80 years, right-handed, and devoid of contraindications for MR scanning. Additional exclusion criteria were employed including incomplete MRI images, poor image quality, and a history of comorbid neurological or psychiatric disease. All participants underwent demographic information collection, clinical assessments, and MRI scanning in one study session. This study was approved by the Institutional Review Board of the Beijing Tiantan Hospital, Capital Medical University, Beijing, China. Written informed consent was obtained from each participant.

### Demographic, Clinical and Neuropsychological Assessment

Demographic and clinical data for both datasets included age, sex, education level, disease duration and the Expanded Disability Status Scale (EDSS) scores. The main dataset had additional clinical scores assessed using the Multiple Sclerosis Functional Composite (MSFC) as well as neuropsychological scores of four cognitive domains assessed using the Brief Repeatable Battery of Neuropsychological Tests (BRB-N) as described previously.^4^ Patients from the main dataset were classified into CIMS and CPMS according to previous studies.^4,26^ Specifically, CIMS patients were defined as those who scored ≥ 1.5 SDs below the control mean on at least 2 subtests of BRB-N, while the others were defined as CPMS. The scores of each of the four cognitive domains were calculated by averaging the scores of all subtests assigned in that domain. The global cognitive function score (global BRB-N) was calculated by averaging the scores of all four cognitive domains.

### MRI Acquisition

#### Main dataset

MRI data were acquired on a 3T MR scanner (General Electric HDx MRI System, GE Medical Devices, Milwaukee, WI) with an 8-channel receive-only head radiofrequency coil. A high-resolution 3D T1-weighted sequence (3DT1) was acquired for identification of T1-hypointense MS lesions, segmentation, and registration (voxel size = 1 mm × 1 mm × 1 mm, echo time [TE] = 3.0 ms, repetition time [TR] = 7.8 ms, matrix size= 256 × 256 × 172, field of view [FOV] = 256 mm × 256 mm, flip angle (FA) = 20°). A T2/proton density–weighted sequence (voxel size = 0.94 mm × 0.94 mm × 4.5 mm, TE = 9.0/80.6 ms, TR = 3,000 ms, FOV = 240 mm × 240 mm, number of slices = 36, FA = 90°) and a fluid-attenuated inversion recovery (FLAIR) sequence (voxel size = 0.86 mm × 0.86 mm × 4.5 mm, TE = 122.3 ms, TR = 9,502 ms, FOV = 220 mm × 220 mm, number of slices = 36, FA = 90°) were acquired for identification and segmentation of T2-hyperintense MS lesions. Resting-state functional MRI (rs-fMRI) was acquired with a T2*-weighted gradient-echo echo-planar imaging sequence (voxel size = 3.4 mm × 3.4 mm × 3 mm, TE = 35 ms, TR = 3,000 ms, matrix size = 64 × 64 × 46, FOV = 220 mm × 220 mm, number of volumes = 100, number of slices = 46, interleaved order). All participants were instructed to relax with their eyes closed during rs-fMRI scanning.

#### Replication dataset

MRI data were acquired on a 3T MR scanner (Philips CX, Best, The Netherlands) including 3DT1, FLAIR, and rs-fMRI. 3DT1 image was acquired using sagittal acquisition with magnetization-prepared rapidly acquired gradient echo (voxel size = 1 mm × 1 mm × 1 mm, echo time [TE] = 3.0 ms, repetition time [TR] = 6.6 ms, matrix size = 196 × 256 × 170, inversion recovering = 880 ms, FA = 8°). FLAIR sequence was acquired using 3D sagittal acquisition with inversion recovering fast spin echo (voxel size = 1 mm × 1 mm × 1 mm, TE = 228 ms, TR = 4,800 ms, inversion time = 1650 ms, FA = 90°). rs-fMRI image was acquired using 2D axial acquisition with field echo EPI (voxel size = 3 mm × 3 mm × 3 mm, TE = 30 ms, TR = 2,000 ms, matrix size= 80 × 80 × 40, number of volumes = 180, number of slices = 40, interleaved order, slice thickness = 3 mm, slice gap = 0.3 mm, FA = 78°) during which all participants were instructed to relax with their eyes closed.

### MRI preprocessing

Lesion filling was performed on the structural 3DT1 images of patients as previously described,^4^ followed by segmentation into grey matter, white matter, and cerebrospinal fluid using SPM12 (v7771) toolbox (http://www.fil.ion.ucl.ac.uk/spm/software/spm12/). The quality of segmentation was assessed manually. rs-fMRI preprocessing was also performed using SPM12. Briefly, individual functional images were first corrected for head motion and acquisition time offsets between slices. No significant differences were found in the maximum and mean frame-wise displacement of head motion between groups in both the main and replication datasets (*p* > 0.05, permutation test). The corrected images were then spatially normalized to the MNI space by applying deformation fields derived from tissue segmentation of structural images. All normalized images further underwent spatial smoothing by a Gaussian kernel with 6-mm full width at half maximum.

### Functional network controllability analysis

We included 400 cortical^27^ and 54 subcortical regions^28^ to calculate functional connectivity networks based on pair-wise Pearson correlation between the time series of all regions. Then, network controllability measures, including average controllability, modal controllability and activation energy, were calculated to quantify how a specific region can influence brain-wide dynamics (Figure 1; See Supplementary Methods for details).^18,19^ Specifically, average controllability measures how easily a brain region move the brain into nearby or easily reachable states, which reflects the brain capacity for low-energy-cost and frequent small adjustments in brain states.^17–19^ Modal controllability, on the contrary, measures a region’s ability to move the brain into difficult or unstable states, which is important for executing high-energy-cost large transitions in brain activity (e.g. between resting and active states).^17–19^ Regional activation energy captures the feasibility or minimum energy required by the given region to induce a transition between brain states.^17–19^ To facilitate interpretation and comparison, region-level controllability measures were averaged into global (whole brain), cortical and subcortical network levels. Cortical network controllability measures were averaged into seven resting-state functional networks,^29^ including the visual network (VN), somatomotor network (SMN), dorsal attention network (DAN), ventral attention network (VAN), limbic network (LN), frontoparietal network (FPN) and default mode network (DMN). Similarly, subcortical network controllability measures were averaged into seven anatomical nuclei, namely the hippocampus, thalamus, amygdala, caudate, nucleus accumbens, putamen and globus pallidus.^28^

**Figure 1.**
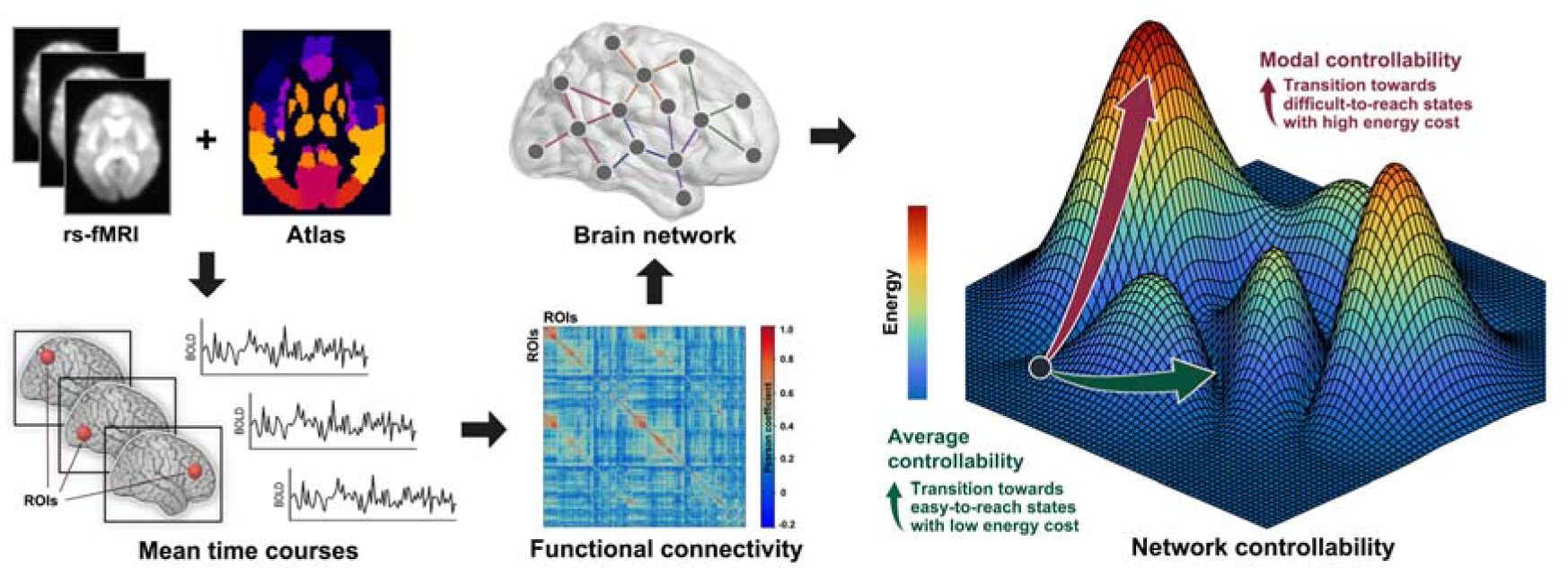
Overview of network controllability calculation. Based on rs-fMRI data from 129 participants and using combined cortical and subcortical parcellations, we extracted regional mean time courses and calculated functional connectivity between pairs of regions to construct brain networks. We then calculated the most commonly used controllability measures to quantify how specific regions influence brain activity transitions throughout the brain. Specifically, average controllability measures how easily a brain region move the brain into nearby or easily reachable states, which reflects the brain capacity for low-energy-cost and frequent small adjustments in brain states. Modal controllability, on the contrary, measures a region’s ability to move the brain into difficult or unstable states, which is important for executing high-energy-cost large transitions in brain activity (e.g. between resting and active states). Regional activation energy captures the feasibility or minimum energy required by the given region to induce a transition between brain states.

### Statistical analysis

Statistical analyses were performed using MATLAB software version R2022b (MathWorks, Inc). A 95% confidence interval was used for each effect. Chi-squared tests were used to compare dichotomous variables (sex). The group comparisons of continuous demographic, clinical, neuropsychological and network controllability variables were performed by permutation tests (10,000 permutations). Age and sex were considered as covariates for neuropsychological and network controllability variables. FDR corrections were performed for multiple comparisons. See Supplementary Methods for details.

#### Classification analysis

We trained linear SVM classifiers to distinguish between MS and controls, as well as between MS with different cognitive status (CIMS vs. CPMS) and between MS with different level of disability (EDSS < 4.0 vs. EDSS ≥ 4.0). Controllability and/or volumetric measures were used as predictive features. A set of performance metrics, including accuracy, precision, sensitivity, specificity, and area under the curve (AUC) were computed to access the performance of the classifiers. See Supplementary Methods for details.

## Results

### Demographic, clinical, and neuropsychological characteristics

#### Main dataset

The MS group was significantly older, had lower education levels, and exhibited worse performance on all subtests of the MSFC as well as all four cognitive domains of the BRB-N than HC group (*p* < 0.05, FDR corrected). Across all patients, 55 were identified as CIMS and 47 as CPMS. CIMS showed worse performance on 9-HPT, PASAT3 and all four cognitive domains than CPMS and HC (*p* < 0.05, FDR corrected). No differences were found between CPMS and HC on cognitive performance (Table 1).

**Table 1.**
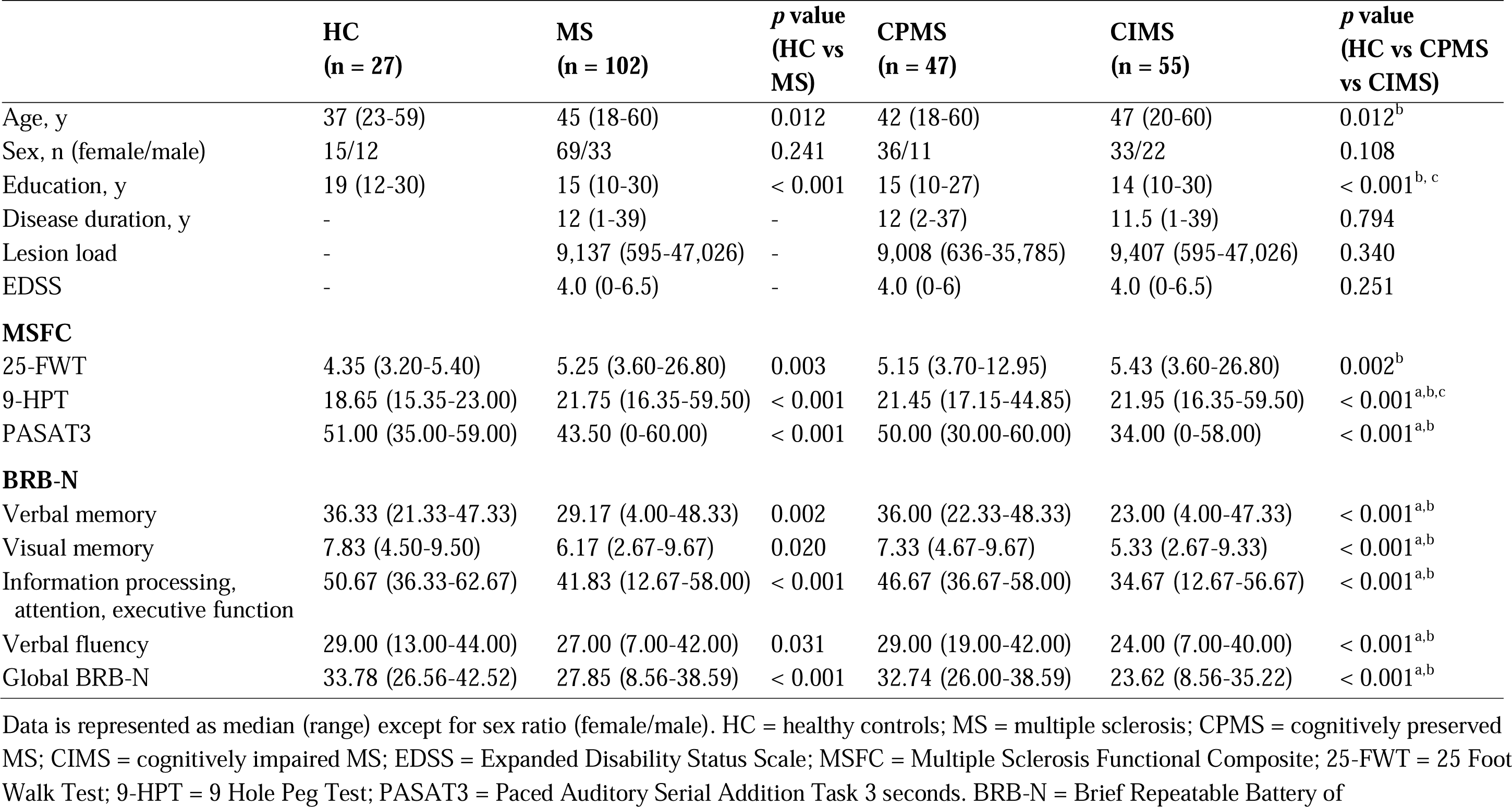
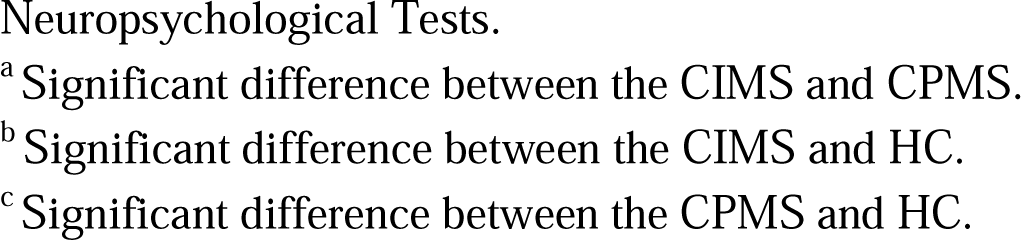
Demographic, clinical, and neuropsychological variables from the main dataset.

|  | **HC**<br>**(n = 27)** | **MS**<br>**(n = 102)** | ***p* value**<br>**(HC vs**<br>**MS)** | **CPMS**<br>**(n = 47)** | **CIMS**<br>**(n = 55)** | ***p* value**<br>**(HC vs CPMS**<br>**vs CIMS)** |
| --- | --- | --- | --- | --- | --- | --- |
| Age, y | 37 (23-59) | 45 (18-60) | 0.012 | 42 (18-60) | 47 (20-60) | 0.012<sup>b</sup> |
| Sex, n (female/male) | 15/12 | 69/33 | 0.241 | 36/11 | 33/22 | 0.108 |
| Education, y | 19 (12-30) | 15 (10-30) | < 0.001 | 15 (10-27) | 14 (10-30) | < 0.001<sup>b, c</sup> |
| Disease duration, y | - | 12 (1-39) | - | 12 (2-37) | 11.5 (1-39) | 0.794 |
| Lesion load | - | 9,137 (595-47,026) | - | 9,008 (636-35,785) | 9,407 (595-47,026) | 0.340 |
| EDSS | - | 4.0 (0-6.5) | - | 4.0 (0-6) | 4.0 (0-6.5) | 0.251 |
| **MSFC** |  |  |  |  |  |  |
| 25-FWT | 4.35 (3.20-5.40) | 5.25 (3.60-26.80) | 0.003 | 5.15 (3.70-12.95) | 5.43 (3.60-26.80) | 0.002<sup>b</sup> |
| 9-HPT | 18.65 (15.35-23.00) | 21.75 (16.35-59.50) | < 0.001 | 21.45 (17.15-44.85) | 21.95 (16.35-59.50) | < 0.001<sup>a, b, c</sup> |
| PASAT3 | 51.00 (35.00-59.00) | 43.50 (0-60.00) | < 0.001 | 50.00 (30.00-60.00) | 34.00 (0-58.00) | < 0.001<sup>a, b</sup> |
| **BRB-N** |  |  |  |  |  |  |
| Verbal memory | 36.33 (21.33-47.33) | 29.17 (4.00-48.33) | 0.002 | 36.00 (22.33-48.33) | 23.00 (4.00-47.33) | < 0.001<sup>a, b</sup> |
| Visual memory | 7.83 (4.50-9.50) | 6.17 (2.67-9.67) | 0.020 | 7.33 (4.67-9.67) | 5.33 (2.67-9.33) | < 0.001<sup>a, b</sup> |
| Information processing,<br>attention, executive function | 50.67 (36.33-62.67) | 41.83 (12.67-58.00) | < 0.001 | 46.67 (36.67-58.00) | 34.67 (12.67-56.67) | < 0.001<sup>a, b</sup> |
| Verbal fluency | 29.00 (13.00-44.00) | 27.00 (7.00-42.00) | 0.031 | 29.00 (19.00-42.00) | 24.00 (7.00-40.00) | < 0.001<sup>a, b</sup> |
| Global BRB-N | 33.78 (26.56-42.52) | 27.85 (8.56-38.59) | < 0.001 | 32.74 (26.00-38.59) | 23.62 (8.56-35.22) | < 0.001<sup>a, b</sup> |

Data is represented as median (range) except for sex ratio (female/male). HC = healthy controls; MS = multiple sclerosis; CPMS = cognitively preserved MS; CIMS = cognitively impaired MS; EDSS = Expanded Disability Status Scale; MSFC = Multiple Sclerosis Functional Composite; 25-FWT = 25 Foot Walk Test; 9-HPT = 9 Hole Peg Test; PASAT3 = Paced Auditory Serial Addition Task 3 seconds. BRB-N = Brief Repeatable Battery of
Neuropsychological Tests.
<sup>a</sup> Significant difference between the CIMS and CPMS.
<sup>b</sup> Significant difference between the CIMS and HC.
<sup>c</sup> Significant difference between the CPMS and HC.

#### Replication dataset

The MS group was significantly younger and had lower education level than HC group (*p* < 0.05, FDR corrected; Supplementary Table S1). Due to the differences in the cognitive assessment scales between the main and replication datasets, further subgroup splitting in terms of CIMS and CPMS were not available on the replication dataset.

### Network controllability changes in MS

#### Main dataset

Significant controllability changes in MS were predominately localized in the subcortical network, particularly in the thalamus. Specifically, at the global (whole brain) level, MS group showed increased average controllability (*p* < 0.001), decreased modal controllability (*p* = 0.008) and decreased activation energy (*p* = 0.016) compared to HC group. When looking at the eight resting-state networks, controllability changes were primarily localized within the subcortical network. Specifically, MS group exhibited increased average controllability (*p* = 0.001), decreased modal controllability (*p* = 0.002) and decreased activation energy (*p* < 0.001) in the subcortical network, compared to HC group (Figure 2). When looked at each of the seven nuclei within the subcortical network separately, the subcortical changes were predominately due to the changes within the thalamus. Specifically, MS group showed increased average controllability (*p* < 0.001), decreased modal controllability (*p* = 0.005) and decreased activation energy (*p* = 0.007) in the thalamus compared to HC (Figure 3). Decreased activation energy was also found in the globus pallidus in MS (*p* = 0.004; Supplementary Figure S1). For the cortical networks, MS-related controllability changes were only found regarding average controllability (VN, SMN, DAN, VAN, FPN and DMN: *p* < 0.05), no differences were observed in any cortical networks regarding modal controllability or activation energy (Supplementary Figure S2).

**Figure 2.**
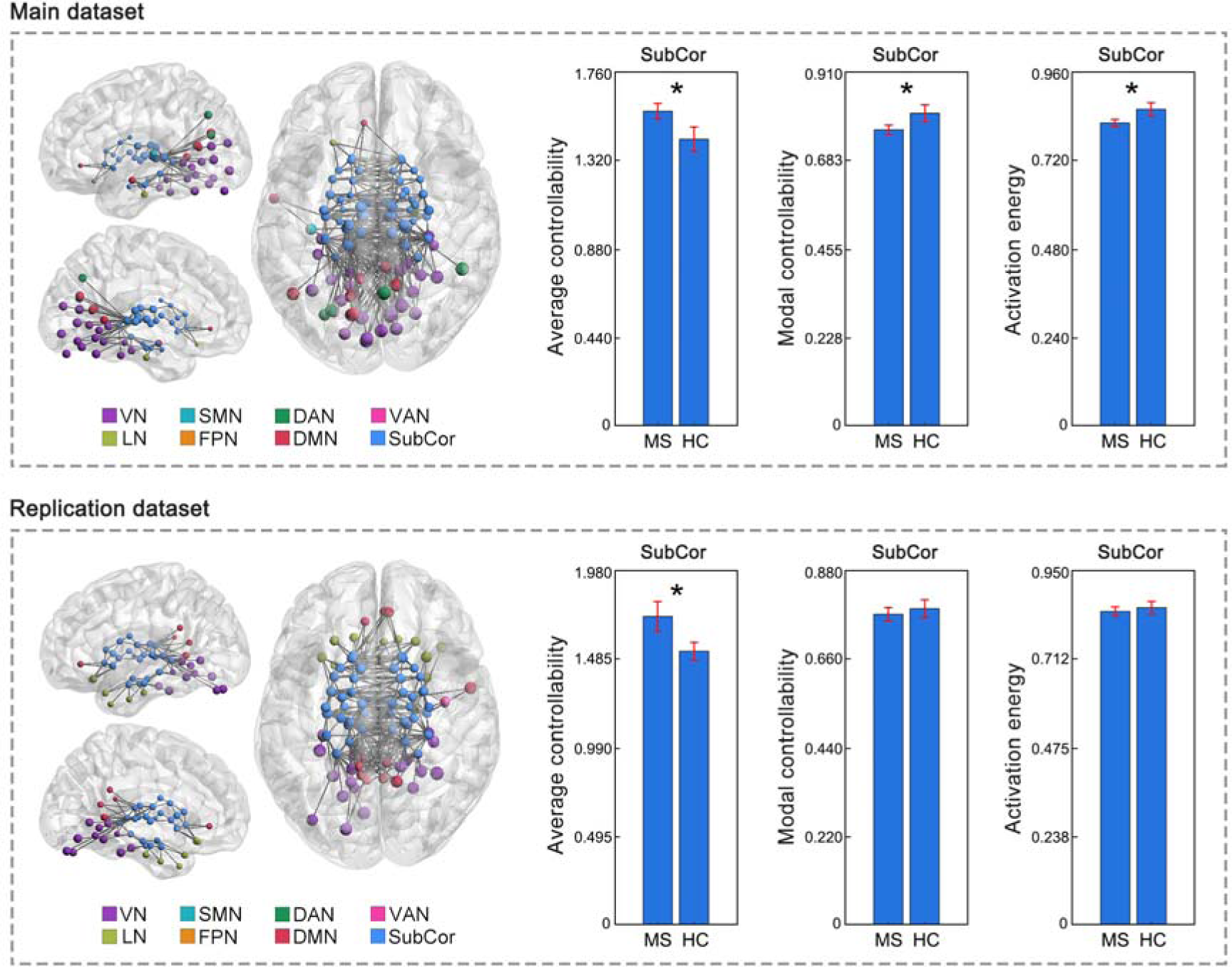
Controllability changes in the subcortical network in MS. Increased average controllability but decreased modal controllability and decreased activation energy in the subcortical network (averaged across the 54 subcortical ROIs) were observed in MS from the main dataset. Replicated increased average controllability in the subcortical network were observed in MS from the replication dataset. Brain network visualizations were generated using BrainNet Viewer^44^ and GRETNA^45^. The nodes and edges illustrate the connections of subcortical regions with other parts of the brain. Specifically, the nodes represent the subcortical regions and the regions to which they are connected. The edges depict the top 200 connections with the highest functional connectivity strength among the whole brain connections. SubCor = subcortical network; \**p* < 0.05, FDR corrected.

**Figure 3.**
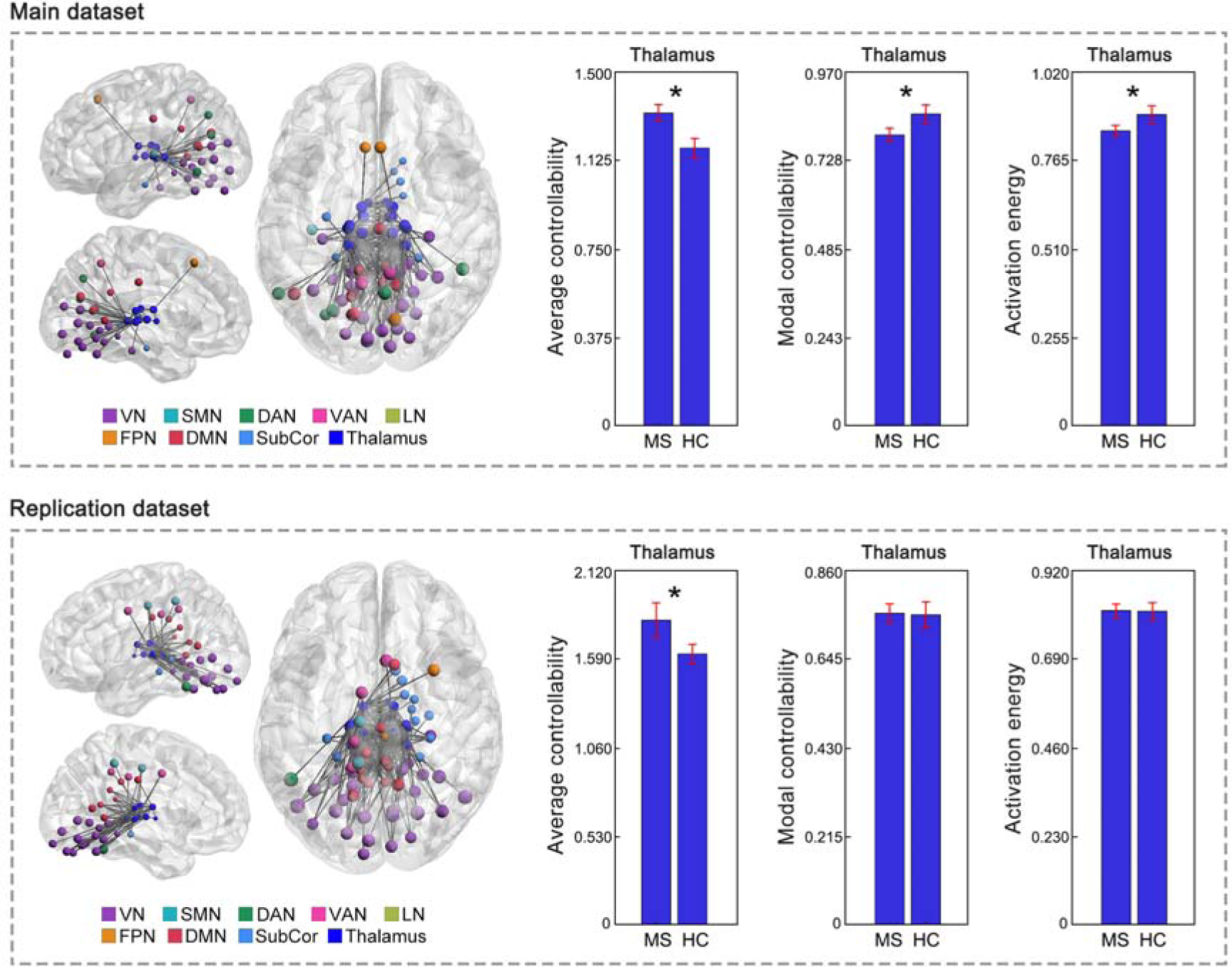
Controllability changes in the thalamus in MS. Increased average controllability but decreased modal controllability and decreased activation energy in the thalamus (averaged across the 16 thalamic ROIs) were observed in MS from the main dataset. Replicated increased average controllability in the thalamus were observed in MS from the replication dataset. Brain network visualizations were generated using BrainNet Viewer^44^ and GRETNA^45^. The nodes and edges illustrate the connections of thalamic regions with other parts of the brain. Specifically, the nodes represent the thalamic regions and the regions to which they are connected. The edges depict the top 200 connections with the highest functional connectivity strength among the whole brain connections. SubCor = subcortical network; \**p* < 0.05, FDR corrected.

#### Replication dataset

Consistent with the main dataset, significant controllability changes in MS were primarily localized in the subcortical network and thalamus in the replication dataset. Specifically, MS exhibited replicated increases in average controllability (*p* = 0.005) in the subcortical network compared to HC (Figure 2). When looked at each of the seven nuclei within the subcortical network separately, MS group showed replicated increases in average controllability (*p* = 0.027) in the thalamus compared to HC (Figure 3). Additionally, increases in average controllability were also observed in other subcortical nuclei, including caudate, nucleus accumbens and putamen (*p* < 0.05, Supplementary Figure S3). No MS-related controllability changes were found in any of the cortical networks.

### Network controllability changes in CIMS and CPMS

Significant controllability differences between CIMS and CPMS were only observed in the thalamus. Specifically, at the whole brain level, both CIMS and CPMS showed increased average controllability (CIMS: p = 0.002; CPMS: p < 0.001), while CIMS additionally exhibited decreased modal controllability (p = 0.009) and decreased activation energy (p = 0.016) compared to HC. When looked at the eight networks, both CIMS and CPMS showed increased average controllability (CIMS: p < 0.001; CPMS: p = 0.008), decreased modal controllability (CIMS: p = 0.006; CPMS: p = 0.008) and decreased activation energy (CIMS: p = 0.007; CPMS: p = 0.003) in the subcortical network compared to HC (Figure 4). When further looked at the seven nuclei within the subcortical network, while both CIMS and CPMS showed increased average controllability in the thalamus (CIMS: p < 0.001; CPMS: p = 0.009), CIMS exhibited significantly higher increases in thalamus than CPMS (p = 0.016, Figure 5). Additionally, only the CIMS group, but not CPMS, showed significant decreases in modal controllability (p = 0.002) and activation energy (p = 0.006) in the thalamus compared to HC (Figure 5). No significant differences were observed between CIMS and CPMS when examining any other parts of the brain aside from the thalamus.

**Figure 4.**
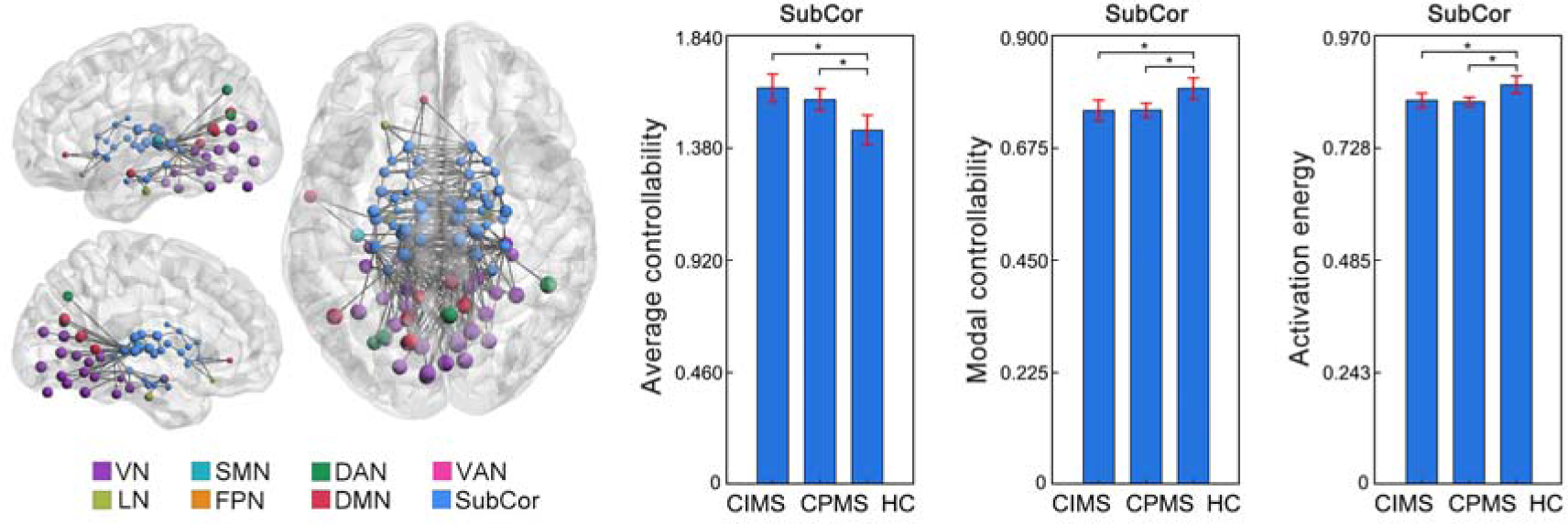
Controllability changes in the subcortical network in CIMS and CPMS. Increased average controllability but decreased modal controllability and decreased activation energy in the subcortical network (averaged across the 54 subcortical ROIs) in both CIMS and CPMS when compared to HC. Brain network visualizations were generated using BrainNet Viewer^44^ and GRETNA^45^. The nodes and edges illustrate the connections of subcortical regions with other parts of the brain. Specifically, the nodes represent the subcortical regions and the regions to which they are connected. The edges depict the top 200 connections with the highest functional connectivity strength among the whole brain connections. SubCor = subcortical network; \**p* < 0.05, FDR corrected.

**Figure 5.**
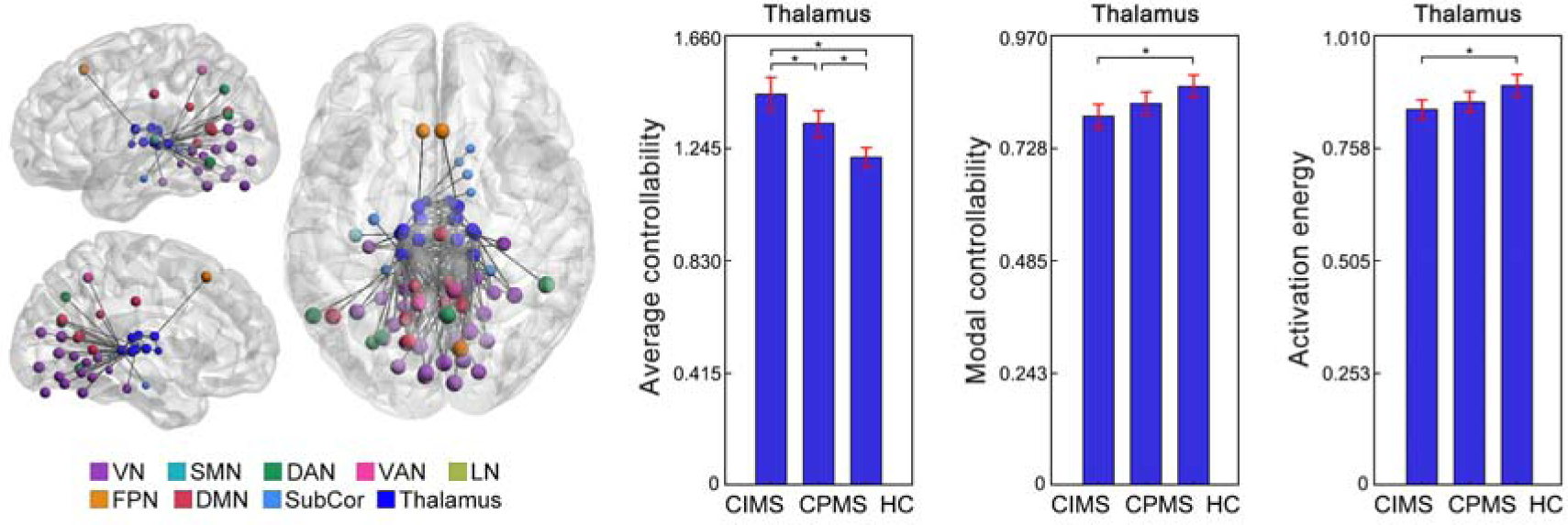
Controllability changes in the thalamus in CIMS and CPMS. Both CIMS and CPMS showed increased average controllability in the thalamus (averaged across the 16 thalamic ROIs) when compared to HC, while CIMS exhibited significantly greater changes than CPMS. Besides, CIMS showed additional decreases in modal controllability and activation energy in the thalamus compared to HC. Brain network visualizations were generated using BrainNet Viewer^44^ and GRETNA^45^. The nodes and edges illustrate the connections of thalamic regions with other parts of the brain. Specifically, the nodes represent the thalamic regions and the regions to which they are connected. The edges depict the top 200 connections with the highest functional connectivity strength among the whole brain connections. SubCor = subcortical network; \**p* < 0.05, FDR corrected.

### Classification performance

Thalamic network controllability measures demonstrated superior classification performance compared to thalamic volume in distinguishing between MS and controls, as well as between MS with different cognitive status and between MS with different level of disability (Figure 6, Table 2). Specifically, classifiers based on both thalamic network controllability and thalamic volume achieved an AUC of 88.3% in distinguishing MS from HC (precision = 93.9%; accuracy = 80.9%; sensitivity = 81.1%; specificity = 80.2%), outperforming those based on thalamic volume alone, which achieved an AUC of 74.5% (precision = 88.8%; accuracy = 68.1%; sensitivity = 68.3%; specificity = 67.6%). Moreover, in differentiating CIMS and CPMS, classifiers based on the thalamic network controllability measures alone achieved an AUC of 80.7% (precision = 75.4%; accuracy = 72.4%; sensitivity = 72.7%; specificity = 72.0%), outperforming those based on the combination of both controllability and volume classifiers (AUC = 68.2%; precision = 64.6%; accuracy = 61.7%; sensitivity = 64.5%; specificity = 58.4%), whereas volumetric classifiers alone failed to differentiate CIMS and CPMS (AUC = 38.2%; precision = 45.3%; accuracy = 41.6%; sensitivity = 41.2%; specificity = 42.1%). Besides, in differentiating MS with different level of disability, classifiers based on the thalamic network controllability measures achieved an AUC of 82.9% (precision = 77.2%; accuracy = 76.3%; sensitivity = 82.4%; specificity = 68.2%), outperforming those based on thalamic volume, which achieved an AUC of 67.8% (precision = 69.3%; accuracy = 64.7%; sensitivity = 67.4%; specificity = 61.2%).

**Figure 6.**
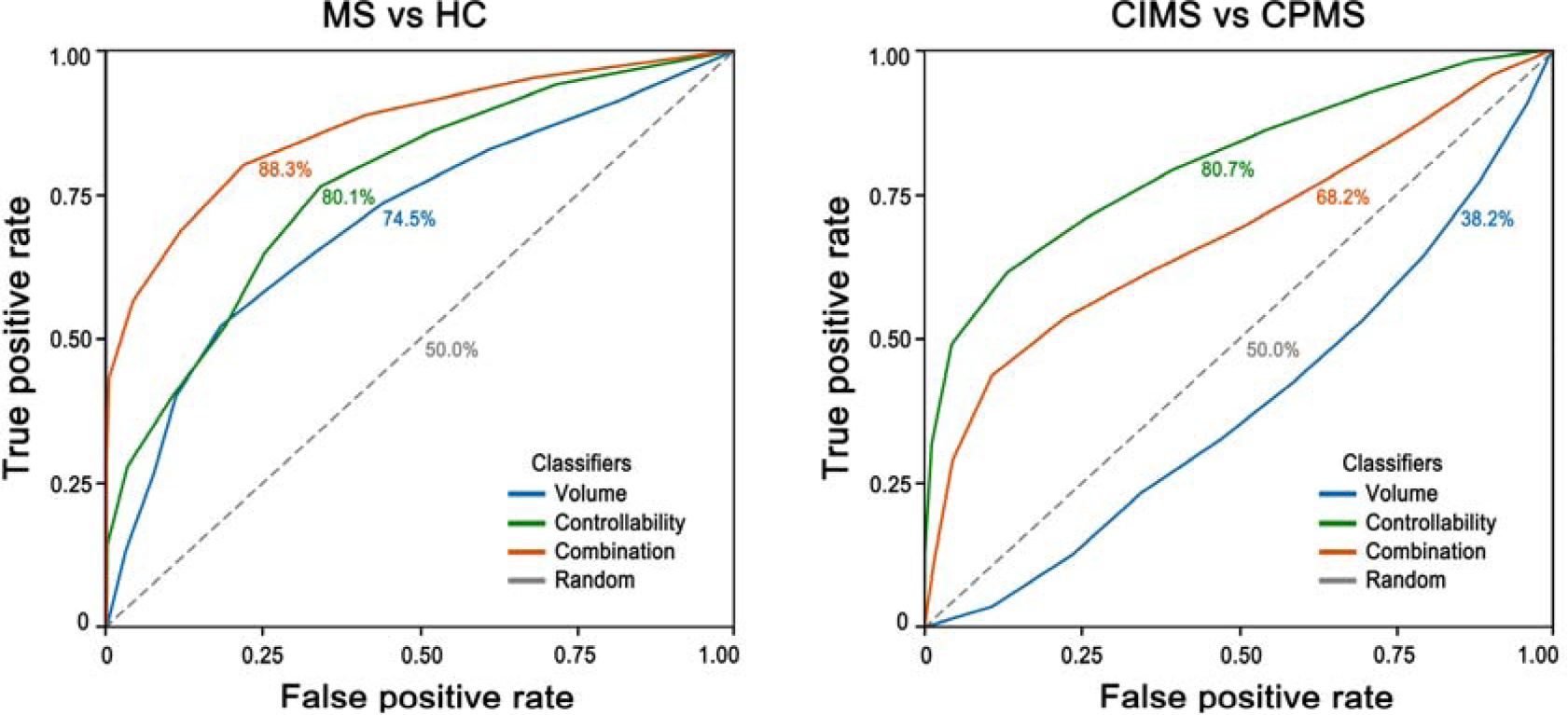
Classification ROC curve derived from volumetric, controllability, combined, and random classifiers. In distinguishing MS from HC, classifiers based on the thalamic network controllability alongside thalamic volume achieved the best performance (AUC = 88.3%) among all the four types of classifiers. In distinguishing CIMS from CPMS, classifiers based on the thalamic network controllability measures alone achieved the best performance (AUC = 80.7%) among all the four types of classifiers. AUC = area under curve; ROC = receiver operating characteristic.

**Table 2.**
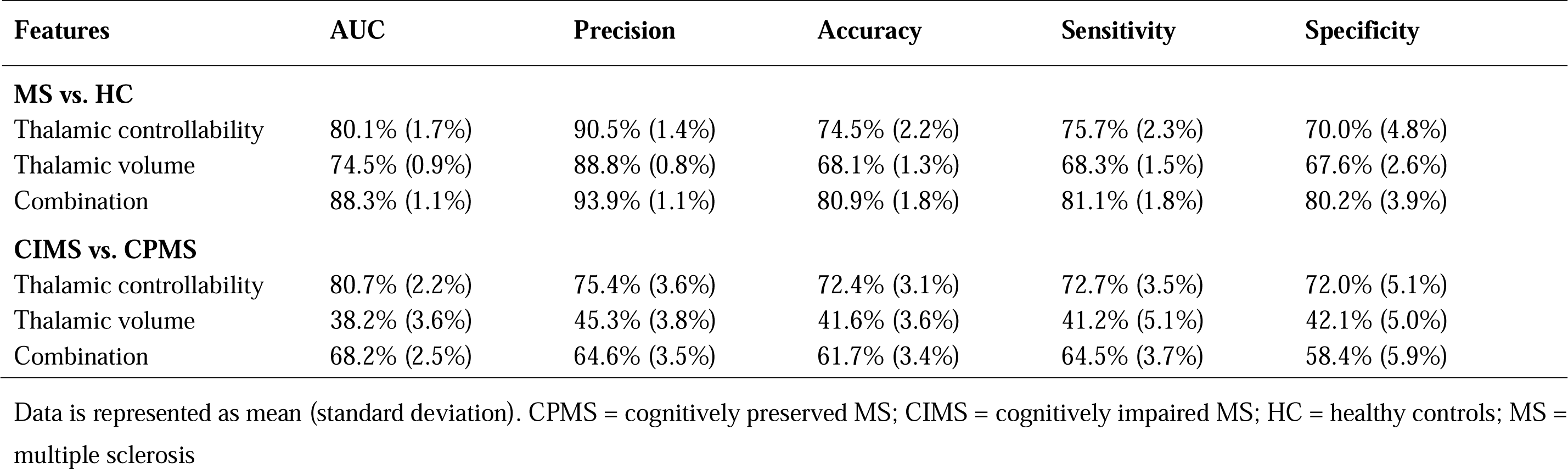
Classification performance using different features.

| Features | AUC | Precision | Accuracy | Sensitivity | Specificity |
| --- | --- | --- | --- | --- | --- |
| **MS vs. HC** |  |  |  |  |  |
| Thalamic controllability | 80.1% (1.7%) | 90.5% (1.4%) | 74.5% (2.2%) | 75.7% (2.3%) | 70.0% (4.8%) |
| Thalamic volume | 74.5% (0.9%) | 88.8% (0.8%) | 68.1% (1.3%) | 68.3% (1.5%) | 67.6% (2.6%) |
| Combination | 88.3% (1.1%) | 93.9% (1.1%) | 80.9% (1.8%) | 81.1% (1.8%) | 80.2% (3.9%) |
| **CIMS vs. CPMS** |  |  |  |  |  |
| Thalamic controllability | 80.7% (2.2%) | 75.4% (3.6%) | 72.4% (3.1%) | 72.7% (3.5%) | 72.0% (5.1%) |
| Thalamic volume | 38.2% (3.6%) | 45.3% (3.8%) | 41.6% (3.6%) | 41.2% (5.1%) | 42.1% (5.0%) |
| Combination | 68.2% (2.5%) | 64.6% (3.5%) | 61.7% (3.4%) | 64.5% (3.7%) | 58.4% (5.9%) |

Data is represented as mean (standard deviation). CPMS = cognitively preserved MS; CIMS = cognitively impaired MS; HC = healthy controls; MS = multiple sclerosis

## Discussion

Our study demonstrated altered functional network control from the thalamus in people with MS, which can be used to predict both MS and cognitive status. Specifically, people with MS showed higher average controllability, lower modal controllability, and lower activation energy in the thalamus. Thalamic network controllability measures proved more effective than thalamic volume alone in distinguishing MS from healthy controls, and in predicting cognitive status. Overall, this study reveals potential mechanisms by which MS-related changes in resting-state functional connectivity impair the thalamus’ ability to drive brain state dynamics, and contribute to cognitive impairment in MS.

The observed changes in network controllability in this work align with previous studies reporting changes in brain activity dynamics in MS.^5,25,30^ The present work, however, extends prior studies by elucidating potential mechanisms through which MS pathology disrupts brain activity dynamics. The controllability measures used in this study provide a holistic view of how specific brain regions drive both subtle and dramatic shifts in cognitive and neural states. Specifically, average controllability measures how easily a brain region move the brain into nearby or easily reachable states, which reflects the brain capacity for low-energy-cost and frequent small adjustments in brain states.^17–19^ Modal controllability, on the contrary, measures a region’s ability to move the brain into difficult or unstable states, which is important for executing high-energy-cost large transitions in brain activity (e.g. between resting and active states).^17–19^ Regional activation energy captures the feasibility or minimum energy required by the given region to induce a transition between brain states.^17–19^ The directions of the observed controllability changes followed a consistent pattern (e.g. increased average controllability, decreased modal controllability, and decreased activation energy), regardless of whether we examined all cortical regions or specific cortical or subcortical nodes. This implies that MS pathology does not induce random changes in brain activity transitions; instead, it produces a disease-related pattern across the brain, consistently characterized by greater difficulties in supporting high-, as opposed to low-energy-cost transitions.

Our results revealed a selective involvement in controllability changes in MS, wherein the subcortical network and particularly the thalamus exhibited the most pronounced changes among the whole brain. A previous controllability study in healthy people found that the subcortical network showed the most imbalance between average controllability and modal controllability - it contributed the most to low-energy-cost state transitions (highest average controllability) while the least to high-energy-cost state transitions (lowest modal controllability).^19^ The healthy cohort in our study conformed to that pattern. However interestingly, MS pathology seems to exacerbate this imbalance in the subcortical network, that is, in MS, the highest average controllability further increased, while the lowest modal controllability further decreased. A recent study in healthy people observed higher average controllability at resting state while higher modal controllability during cognitive tasks.^31^ This was interpreted as the resting state being a ‘ground state’ that maintains the energy cost at a relatively low level, whereas performing cognitive tasks represents an ‘excited state’ that consumes a large amount of energy to facilitate cognitive functions.^31^ Given that the subcortical network, particularly the thalamus, is one of the most vulnerable brain areas affected by MS,^6–8,10–15,32–34^ the controllability imbalance might be a manifestation of disease attacks in these areas. This provides a potential mechanism explaining increased cognitive impairment with disease progression. The thalamus increasingly ‘drives’ the brain transitions towards low-energy-cost states (like the resting state) while relinquishing transitions towards high-energy-cost states (like the cognitive task state). This process might help preserve fundamental neural activities against disease attacks but, on the other hand, impact brain functioning associated with high-energy-cost activities, typically cognitive processes.

The analysis of cognitive subgroups further supported this view that thalamic controllability changes may impact cognitive performance in patients. While both CIMS and CPMS showed controllability changes compared to HC, CIMS exhibited significantly greater changes in the thalamus than CPMS. This indicated that CIMS experiences greater difficulty in the thalamus supporting brain transitions towards difficult-to-reach states, which are typically associated with high-energy-cost cognitive functions.^17–19,31^ As a key hub with strong connections across the brain and with known roles in cognition (e.g. the anterior thalamic nuclei is critical for memory), thalamic damage has received considerable attention in MS as a predictor of cognitive impairment.^6–8,34^ Several studies converge to show that the thalamus is an early site of pathology in MS,^11–13^ while its structure and function have been reported to determine the severity of cognitive impairment in patients.^7,35^ These findings strongly suggest thalamic changes as promising biomarkers for cognitive dysfunction in MS. However, a key question would be: how does MS pathology progress from only localized damage in thalamus (and a few neighbouring regions) in the early disease phase^11–16,36^ towards the widespread ‘network collapse’ across the brain in the late disease phase,^16,36,37^ culminating in cognitive dysfunction. The present study considering control processes from a bioengineering perspective sheds lights on this. Recent evidence showed that CIMS, compared to CPMS, required more control energy to transition between brain states.^25^ In light of this evidence, our findings suggest that early disease attacks produce a shift in controllability in the thalamus that promotes brain network changes towards low-energy-cost activity patterns, which would preserve energy for essential neural activities. However, this ‘energy-saving’ mode restricts the brain’s efficiency to support high-energy-cost cognitive functions in patients, gradually leading to impaired performance in cognitive functions. This highlights a potential pathobiological mechanism linking thalamic changes to cognitive impairment in MS.

Our results demonstrate that thalamic network controllability can help distinguish MS patients from healthy controls and differentiate between patients with different level of disability and cognitive impairment, with better performance compared to previously reported classification analysis in this disease.^38^ While both thalamic volume and controllability contribute to classifying patients with different levels of disability, only thalamic controllability achieves high classification performance in distinguishing between CIMS and CPMS. These findings suggest that thalamic controllability offers complementary information to volumetric measures in distinguishing patient groups. In particular, controllability appears to be more specifically associated with cognitive impairment, rather than just reflecting a general marker of network collapse. Importantly, the mechanistic interpretation of controllability changes potentially allows for explainable diagnosis. This is a key advantage compared to typical diagnostic tools based on statistical classification, which only allows for a yes or no answer.

Notably, a recent longitudinal study in MS reported that volume changes in deep grey matter regions, including the thalamus, were correlated with worsening clinical disability over the course of the disease.^32^ In parallel, altered functional dynamics in the subcortical areas, particularly a hyperflexible reorganization of brain activity, have also been observed in MS, showing significant associations with the development of clinical impairment.^39^ These findings underscore the potential importance of the dynamic profiles of subcortical regions, like the thalamus, in understanding the progression of clinical dysfunctions in MS. Emerging evidence highlights a complex and dynamic interplay between cognitive impairment, physical disability progression, and thalamic structural and functional changes in MS. Longitudinal studies have demonstrated that cognitive impairment at diagnosis can serve as a predictor of subsequent disability milestones, including faster physical decline and poorer clinical outcomes.^40–42^ Likewise, thalamic structural and functional changes at baseline has been shown to predict not only future cognitive dysfunctions but also worsening disability, underlining the thalamus as a central region implicated in broader neurodegenerative processes in MS.^6–8^ Conversely, individuals with more severe disability tend to exhibit greater degrees of cognitive dysfunction and more pronounced thalamic alterations.^43^ These findings suggest that cognitive impairment, thalamic changes, and clinical disability may not occur in isolation but rather reflect shared pathological pathways or processes. Taken together, the above evidence supports the hypothesis that changes in thalamic controllability may serve as an early biomarker of cognitive decline and potentially contribute to the progression of cognitive dysfunction in MS. Future longitudinal studies tracking network controllability alongside cognitive outcomes in MS will be essential to formally test this hypothesis.

This study has several limitations. First, although the CIMS and CPMS groups did not differ significantly in age or sex, the overall MS group was older than the healthy control group. To mitigate the potential influence of these demographic differences on our findings, we included age and sex as covariates in all relevant statistical models. Second, the imaging scanning parameters and cognitive assessments differed between the two independent datasets. However, the observed similar pattern of results demonstrates the robustness of the findings despite these differences in acquisition and scanner between datasets. Future studies with multiple MS cohorts using paired scanning parameters and cognitive assessments are warranted to better replicate the controllability changes in patients with different cognitive statuses. Third, our study is cross-sectional; further longitudinal studies are needed to determine how network controllability develops with disease progression in MS.

## Conclusion

This study moves beyond confirming thalamic changes in MS by characterizing how the MS-related damage significantly impact the thalamus ability to drive brain-wide dynamic activity, and how this helps explain cognitive impairment in this condition. Our results demonstrate that the thalamus in cognitively impaired MS patients is less able to facilitate brain transitions crucial for high-energy-cost cognitive functions, providing novel insights into the pathological mechanisms linking thalamic functional changes to cognitive impairment in MS.

## Data and Code Availability

The data and code that support the results of this study are available from the corresponding author upon reasonable request and with permission.

## Author Contributions

Y.L., N.T.B., and N.M. contributed to the conception and design of the study. I.L., Y.L., N.T.B., and N.M. contributed to the acquisition of data. Y.Y., A.W., Z.Z., M.C.L., N.T.B., and N.M. contributed to analysis or interpretation of data. Y.Y., A.W., V.T., Y.L., N.T.B., and N.M. contributed to drafting/revision of the manuscript.

## Acknowledgements

This work was funded by research grants from the MS Society UK and MRC UK (MR/X005267/1). We extend our gratitude to Dr. Danka Jandric, Dr. Elizabeth McManus, Jessica Haigh, Megan Sheppard, Katie Moran, Rose-Marie Kouwenhoven, and all our colleagues from the MISC and NIMO labs for their kind support throughout this study.

## Supplementary Material

Supplementary materials are available online.

## Abbreviations

AUC: Area under curve
BRB-N: Brief Repeatable Battery of Neuropsychological Tests
CIMS: cognitively impaired multiple sclerosis
CPMS: cognitively preserved multiple sclerosis
DAN: dorsal attention network
DMN: default mode network
EDSS: Expanded Disability Status Scale
FA: flip angle
FLAIR: fluid-attenuated inversion recovery
FPN: frontoparietal network
HC: healthy control
LN: limbic network
MS: multiple sclerosis
MSFC: Multiple Sclerosis Functional Composite
PASAT3: Paced Auditory Serial Addition Task 3 seconds
ROC: receiver operating characteristic
rs-fMRI: resting-state functional MRI
SD: standard deviation
SMN: somatomotor network
VAN: ventral attention network
VN: visual network
25-FWT: 25 Foot Walk Test
3DT1: 3D T1-weighted sequence
9-HPT: 9 Hole Peg Test.

