## Supplementary Materials for "Thalamic Network Controllability Predicts Cognitive Impairment in Multiple Sclerosis"

### Supplementary Methods

#### **Functional network construction**

The functional network was constructed based on a combined parcellation scheme of 454 regions of interest (ROIs), including 400 cortical regions defined by the Schaefer atlas according to the functional network organization of cortical areas^1^ and 54 subcortical regions defined by a subcortical parcellation derived from functional connectivity gradients of the subcortical areas.^2^ We extracted and averaged the time series of each voxel within each ROI, and then calculated the Pearson correlation coefficient of the time series between each ROI pair to characterize functional connectivity. Non-significant connections (*p* ≥ 0.05) were excluded, and only those connections with neurophysiological significance (*p* < 0.05) were preserved in the functional networks.

#### **Network controllability calculation**

Following previous studies^3–5^, we constructed a simplified linear time-invariant control system to characterize the controllability of functional brain networks. Specifically, we described the dynamics of brain state transitions as a linear time-invariant dynamical model of the form:

$$\dot{x}\left( t \right)=Ax\left( t \right)+Bu(t)$$

where $x\left( t \right)$ represents the brain state at time $t$; $A$ is the time-invariant system matrix modelling the interactions between different nodes of the brain network and was defined as a normalized version of the functional connectivity matrix; B is a matrix comprising columns of the identity matrix indicating the set of network nodes used to control the network; $u(t)$ denotes the input control energy injected at each control node at time $t$. To calculate $A$, we used a modified Laplacian normalization of the functional connectivity matrix $F$ to ensure the stability of the dynamic system.^5^ Specifically, we calculated the Laplacian matrix as:

$$L_{ij}= \delta_{ij}\sum_{k=1}^{K} \left| F_{iK} \right|-F_{ij}$$

where $F_{ij}$ is the $(i,j)$ element of the functional matrix; $K$ is the total number of nodes in the network; $\delta_{ij}$ is the Kronecker delta ($\delta_{ij}=1$ if $i=j$ and $\delta_{ij}=0$ otherwise). The functional connectivity matrix was then normalized into a system matrix as:

$$A= \frac{-L}{\lambda_{max}(L)}$$

with $\lambda_{max}(L)$ denoting the maximum absolute eigenvalue of the Laplacian matrix.

To characterize the ability of a given brain region (a network node) to control the whole brain network, we calculated the most widely used controllability measures for each region (network node), including average controllability, modal controllability and activation energy (Figure 1).^3,5^ Specifically, average controllability measures how easily a brain region move the brain into nearby or easily reachable states, which reflects the brain capacity for low-energy-cost and frequent small adjustments in brain states.^3,5,6^ Modal controllability, on the contrary, measures a region’s ability to move the brain into difficult or unstable states, which is important for executing high-energy-cost large transitions in brain activity (e.g. between resting and active states).^3,5,6^ Regional activation energy captures the feasibility or minimum energy required by the given region to induce a transition between brain states.^3,5,6^ Average controllability of region $i$ was calculated as the trace of the controllability Gramian matrix $W_{i}$, which is defined as

$$W_{i}=\int_{0}^{\infty} e^{At}B_{i}B_{i}^{T}e^{A^{T}t}dt$$

where $e^{At}$ and $e^{A^{T}t}$ denote the matrix exponential of the matrices $At$ and $A^{T}t$, respectively; $B_{i}$ is the$i$-th column of the identity matrix, indicating that region $i$ is used as control node. Modal controllability of region $i$ was calculated as

$$\phi_{i}=\sum_{j=1}^{N} (1-\lambda_{j}^{2}(A))v_{ij}^{2}$$

with $\lambda_{j}\left( A \right)$ representing the $j$-th eigenvalue of the system matrix $A$, and $V=\left[ v_{ij} \right]$ denoting the eigenvector matrix of $A$. Activation energy of region $i$ was defined as

$$\varepsilon_{i}=\frac{1}{2}d_{i}^{T}W^{-1}d_{i}$$

where $d_{i}$ is a column vector with the $i$-th element equal to one and the others being zero, indicating the activation of the $i$-th region.

#### **Statistical analysis**

Statistical analyses were performed using MATLAB software version R2022b (MathWorks, Inc). A 95% confidence interval was used for each effect. Chi-squared tests were used to compare dichotomous variables (sex). Group comparisons of continuous demographic, clinical, neuropsychological, and network controllability variables were conducted using non-parametric permutation tests with 10,000 permutations. Age and sex were considered as covariates for neuropsychological and network controllability variables. An additional model that also included education level as a covariate was also tested. Specifically, we first computed the test statistic (T or F) using two-sample t-tests (e.g., for disease duration and EDSS), ANOVA (e.g., for age), or ANCOVA (e.g., for education, MSFC, BRB-N, volume, and controllability measures). We then generated an empirical null distribution of the test statistic by randomly permuting group labels and recalculating the statistic 10,000 times. The empirical p-value was calculated as the proportion of permuted test statistics that were equal to or exceeded the absolute value of the observed statistic. To correct for multiple comparisons, we applied false discovery rate (FDR) correction at *q* < 0.05. For variables showing significant group differences across the three groups, post hoc pairwise comparisons were also performed using permutation t-tests (10,000 permutations) following the same procedure.

**Assessment of controllability changes in MS**

The differences on global, cortical, and subcortical controllability variables between MS and HC groups were identified by permutation tests (10,000 times, FDR corrected). Age and sex were considered as covariates. Notably, controllability values that exceeded 1.5 times the interquartile range above quartile 3 or fell below 1.5 times the interquartile range beneath quartile 1 within each group were identified as outliers and then excluded from this analysis (Supplementary Table S2 and S3). To assess the robustness of our findings, we performed sensitivity analyses by (1) replacing the identified outliers with their nearest non-outlier sample and (2) conducting the analysis without any outlier removal or replacement. To test the reliability of our results, we further assessed controllability changes in MS in the replication dataset. All analysis procedures were performed identically between the two datasets.

**Assessment of controllability changes in CIMS and CPMS**

To explore how controllability changes reflect cognitive impairment in MS, we further assessed controllability changes in CIMS and CPMS. Specifically, permutation tests (10,000 times, FDR corrected) were first applied between CIMS, CPMS and HC. For those controllability measures that showed significant differences among the three groups, *post hoc* tests were further used for the pairwise comparisons between each two groups. Again, age and sex were considered as covariates. Outliers of controllability measures were handled as described before.

#### **Classification analysis**

Classification analyses were performed between MS and HC as well as between CIMS and CPMS groups. Specifically, we trained linear SVM binary classifiers for each group pair, using controllability measures and/or volumetric measures as predictive features. Out-of-sample classification performance was evaluated using 10-fold cross-validation, during which stratification was implemented to ensure representative group proportions within each fold. Prior to model training, both the training and testing datasets for each fold underwent standardization based on the mean and SD of the training dataset. Linear SVM classifiers were then trained on the standardized training dataset, and then applied to the testing dataset. After training, a set of performance metrics including accuracy, precision, sensitivity, specificity and area under the curve (AUC) were computed to characterize the performance of the classifiers. This classification procedure was iterated 100 times to increase reliability. The resulting mean values of each performance metric across all repetitions were reported to provide a robust estimation of classification performance. Notably, the effects of imbalance between classes were taken into consideration by assigning weights to each class according to the number of samples in that class.

### Supplementary Results

#### **Sensitivity analyses of outlier-handling approach**

The main results remained consistent across different outlier-handling approaches, no matter by removing (*p_rem_*), replacing (*p_rep_*), or reserving outliers (*p_res_*). Specifically, when comparing between MS and HC, MS showed increased average controllability (*p_rem_* < 0.001, *p_rep_* < 0.001, *p_res_* = 0.008), decreased modal controllability (*p_rem_* = 0.005, *p_rep_* = 0.005, *p_res_* = 0.005) and decreased activation energy (*p_rem_* = 0.007, *p_rep_* = 0.007, *p_res_* = 0.007) in the thalamus compared to HC. When comparing between CIMS, CPMS and HC, while both CIMS and CPMS showed increased average controllability in the thalamus (CIMS: *p_rem_* < 0.001, *p_rep_* < 0.001, *p_res_* = 0.008; CPMS: *p_rem_* = 0.018, *p_rep_* = 0.009, *p_res_* = 0.026), CIMS exhibited significantly higher increases in thalamus than CPMS (*p_rem_* = 0.018, *p_rep_* = 0.017, *p_res_* = 0.022). Additionally, only the CIMS group, but not CPMS, showed significant decreases in modal controllability (*p_rem_* = 0.002, *p_rep_* = 0.002, *p_res_* = 0.002) and activation energy (*p_rem_* 0.006, *p_rep_* = 0.006, *p_res_* = 0.006) in the thalamus compared to HC. No significant differences were observed between CIMS and CPMS when examining any other parts of the brain aside from the thalamus.

#### **Sensitivity analysis of the effect of education level**

The main results remained consistent, whether we included only age and sex as covariates (*p_AgeSex_*), or also included education level as a covariate alongside age and sex (*p_AgeSexEdu_*) in our statistical models. Specifically, MS group showed increased average controllability (*p_AgeSex_* < 0.001, *p_AgeSexEdu_* < 0.001), decreased modal controllability (*p_AgeSex_* = 0.005, *p_AgeSexEdu_* = 0.007) and decreased activation energy (*p_AgeSex_* = 0.007, *p_AgeSexEdu_* = 0.008) in the thalamus compared to HC. For comparison between cognitive subgroups, while both CIMS and CPMS showed increased average controllability in the thalamus (CIMS: *p_AgeSex_* < 0.001, *p_AgeSexEdu_* < 0.001; CPMS: *p_AgeSex_* = 0.009, *p_AgeSexEdu_* = 0.008), CIMS exhibited significantly higher increases in thalamus than CPMS (*p_AgeSex_* = 0.01, *p_AgeSexEdu_* = 0.020). Additionally, only the CIMS group, but not CPMS, showed significant decreases in modal controllability (*p_AgeSex_* = 0.002, *p_AgeSexEdu_* = 0.003) and activation energy (*p_AgeSex_* = 0.006, *p_AgeSexEdu_* = 0.005) in the thalamus compared to HC.

**Supplementary Table S1.** Demographic and clinical variables from the replication dataset

|  | **HC (n = 45)** | **MS (n = 95)** | ***p* value (HC vs MS)** |
| --- | --- | --- | --- |
| Age, y | 35 (22-77) | 33 (17-80) | 0.014 |
| Sex, n (female/male) | 32/13 | 65/30 | 0.747 |
| Education, y | 16 (5-19) | 15 (5-19) | 0.004 |
| Disease duration, y | - | 3 (0.03-36) | - |
| EDSS | - | 2.5 (0-8) | - |

Data are represented as median (range) except for sex ratio (female/male). Abbreviations: HC = healthy controls; MS = multiple sclerosis; EDSS = Expanded Disability Status Scale.

**Supplementary Table S2.** Numbers of outliers when comparing between MS and HC

| **Measures** | **MS** | **HC** |
| --- | --- | --- |
| Average controllability | 7 | 4 |
| Modal controllability | 0 | 0 |
| Activation energy | 0 | 0 |

HC = healthy controls; MS = multiple sclerosis.

**Supplementary Table S3.** Numbers of outliers when comparing CIMS, CPMS and HC

| **Measures** | **CIMS** | **CPMS** | **HC** |
| --- | --- | --- | --- |
| Average controllability | 2 | 3 | 4 |
| Modal controllability | 0 | 2 | 0 |
| Activation energy | 0 | 0 | 0 |

MS = multiple sclerosis. CPMS = cognitively preserved MS; CIMS = cognitively impaired MS.

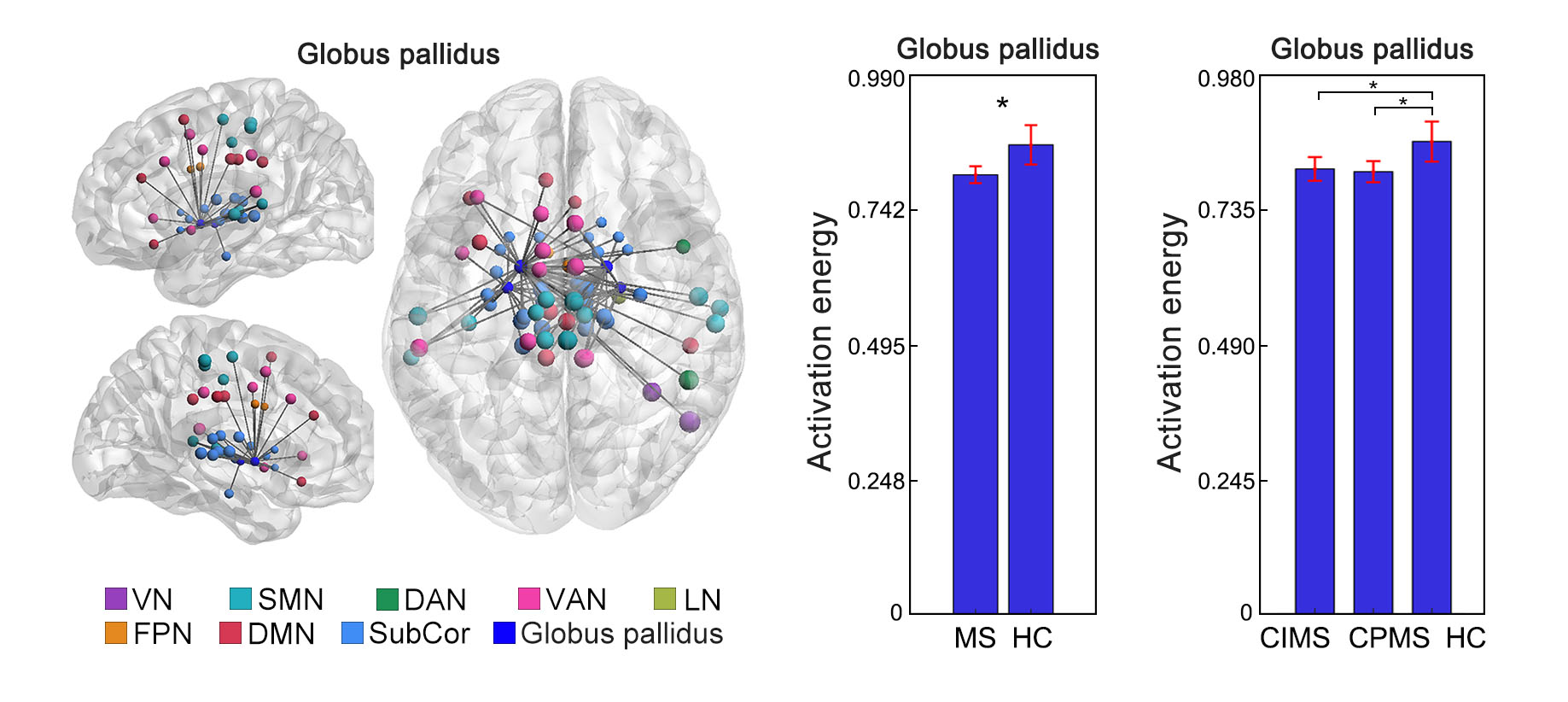

**Supplementary Figure S1**. **Additional controllability changes in subcortical nuclei in MS from the main dataset.** Decreased activation energy in the globus pallidus was observed in MS compared to HC. Both CIMS and CPMS showed the decreases compared to HC. No difference was found between CIMS and CPMS.
CIMS = cognitively impaired multiple sclerosis; CPMS = cognitively preserved multiple sclerosis; HC = healthy control; MS = multiple sclerosis; ^*^*p* < 0.05, FDR corrected.

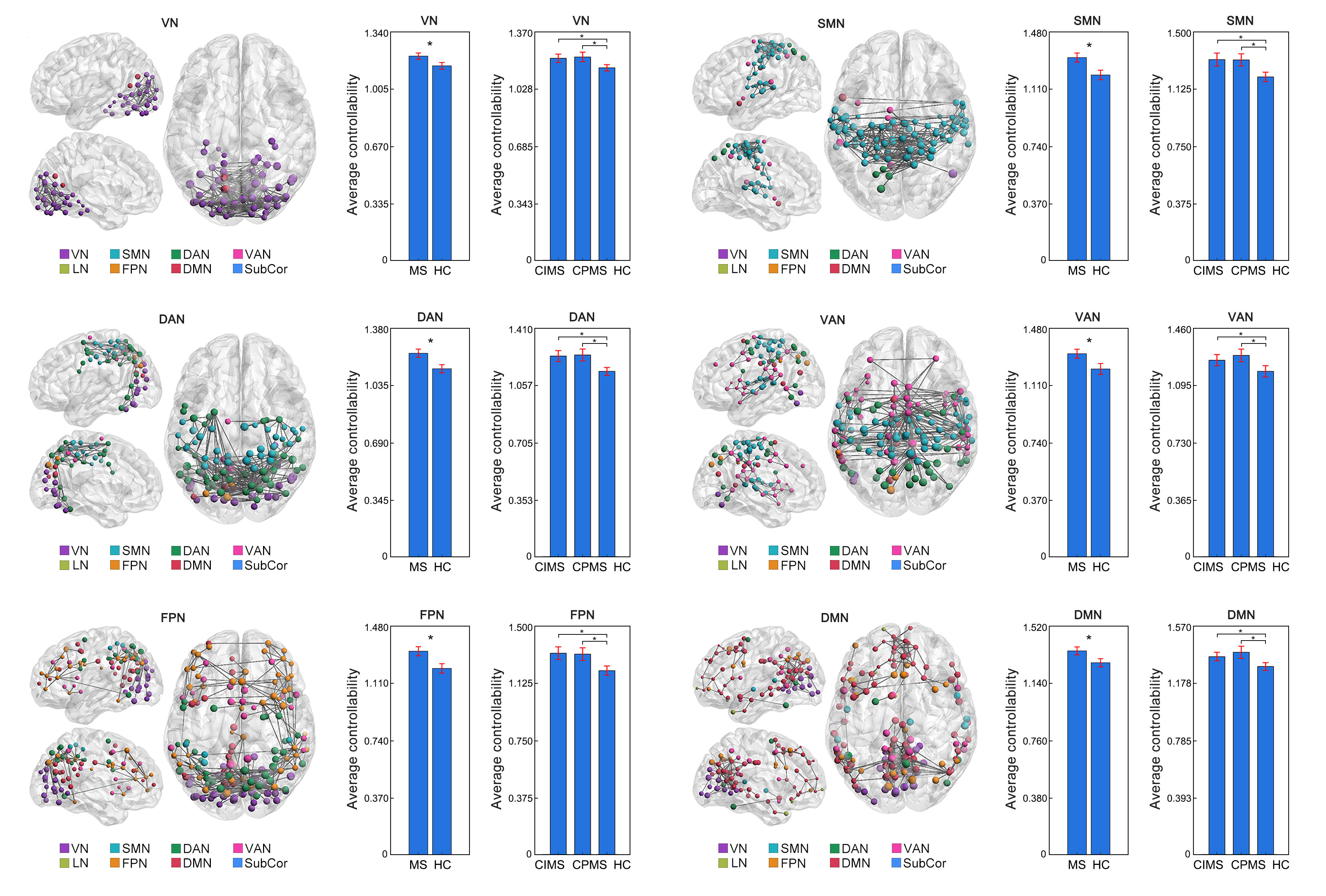

**Supplementary Figure S2. Additional controllability changes in the cortical networks in MS from the main dataset.** Increased average controllability was found in the VN, SMN, DAN, VAN, FPN, and DMN in MS when compared to HC. Both CIMS and CPMS showed increases compared to HC. No difference was found between CIMS and CPMS.
CIMS = cognitively impaired multiple sclerosis; CPMS = cognitively preserved multiple sclerosis; DAN = dorsal attention network; DMN = default mode network; FPN = frontoparietal network; HC = healthy control; LN = limbic network; MS = multiple sclerosis; SMN = somatomotor network; SubCor = subcortical network; VAN = ventral attention network; VN = visual network; ^*^*p* < 0.05, FDR corrected.

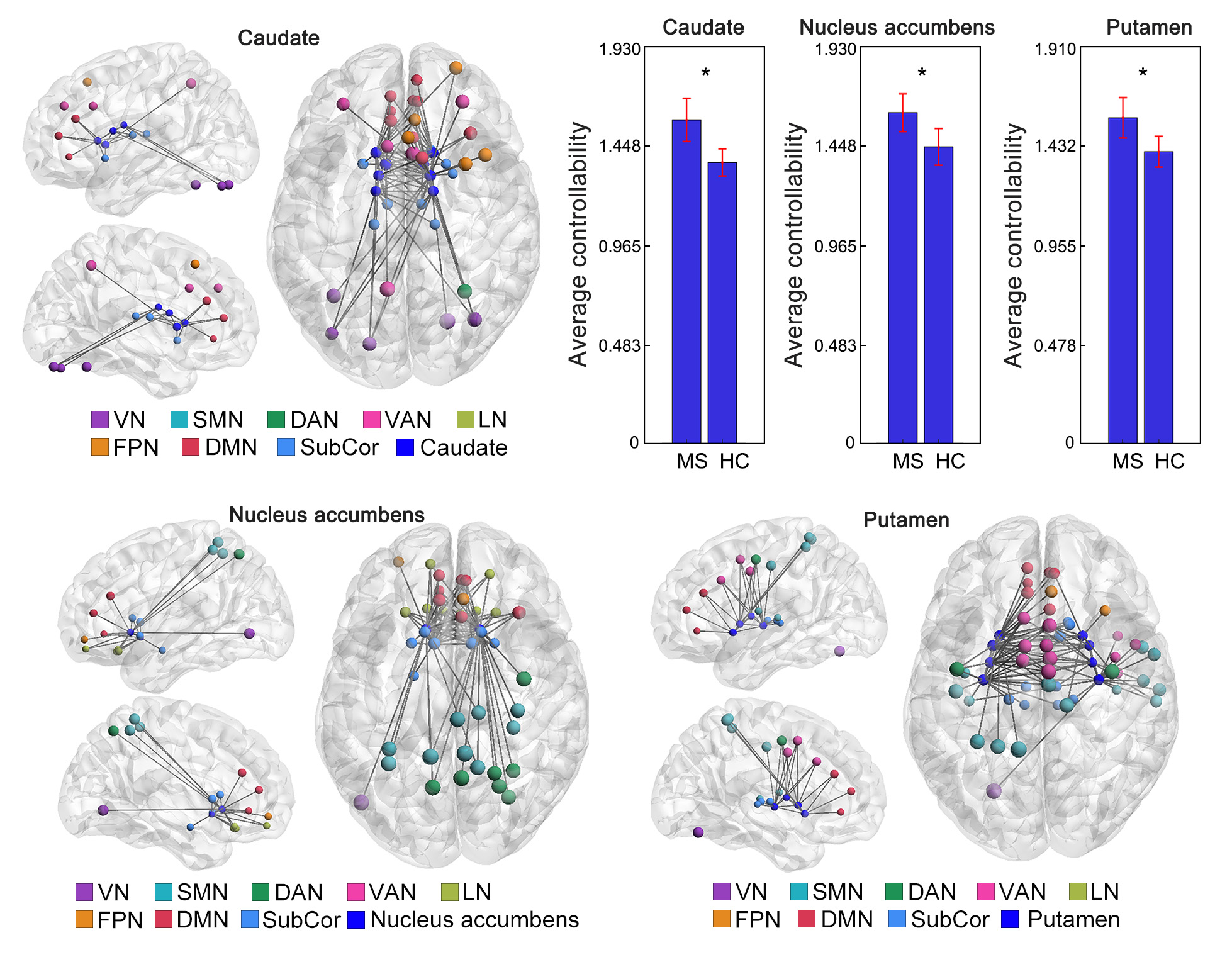

**Supplementary Figure S3. Additional controllability changes in subcortical nuclei in MS from the replication dataset.** Additional increased average controllability was found in the caudate, nucleus accumbens and putamen in MS from the replication dataset when compared to HC.
HC = healthy control; MS = multiple sclerosis; ^*^*p* < 0.05, FDR corrected.
